# Estimation of the biosorption potential of certain representatives of the genus Bacillus in interaction with lead cation *in vitro*

**DOI:** 10.1101/2021.12.27.474245

**Authors:** A.N. Sizentsov, L.V. Galaktionova, O.K. Davydova, A.N. Nikiyan, Ya.A. Sizentsov

## Abstract

The article presents data on the physicochemical and metabolically dependent mechanisms of detoxification by microorganisms of heavy metals that enter the environment during anthropogenic pollution. The taxonomic and physiological-biochemical diversity of microorganisms capable of neutralizing toxicants has been demonstrated. In the experimental part of the study, the combination of the methods used made it possible to fully assess the degree of toxicity and the effect of lead cations on the growth of bacteria of the genus Bacillus in a model experiment. Thus, the use of atomic absorption spectrophotometry and atomic force microscopy gave an idea of the level of biosorption of a given xenobiotic element from a substrate with localization of inactive forms of lead. The data obtained indicate the presence of an inhibitory effect of Pb(NO_3_)_2_ and Pb(CH_3_COO)_2_ in concentrations from 1 mM to 0.063 mM about the test organisms under study. The presence of cations with a nutrient substrate gives a false-positive idea of the degree of influence of lead on the growth of the studied microorganisms, since an increase in optical density in test samples is due to high sorption characteristics and, as a consequence, is characterized by an increase in relative optical density. An assessment of the detoxification mechanisms, which is expressed by active bioaccumulation of lead on the surface elements of the microbial cell up to 65% at the point of entering the stationary growth phase, indicates the promising use of representatives of this group of microorganisms as microbial bioremediation and correctors of excess content of this element in the body or ecosystem as a whole.

## INTRODUCTION

The study of individual links of geochemical cycles in the biosphere and the biological role of metals in ecosystems suggests a dual nature. A large list of metals is necessary for the normal course of the vital processes of living organisms, however, with an increase in their content in the ecosystem, they can hurt the vital activity of organisms and demonstrate high toxicity [1–6]. The boundary between inert, biological, and toxic concentrations of metals is unclear, which complicates the study of their impact on the environment. The amount at which some metals become toxic depends not only on their concentration in the components of the ecosystem but also on the characteristics of the biochemical cycle and the chemical properties of the element in a specific geochemical setting [7–10].

Natural sources of heavy metals are emitted from volcanoes, transport of continental dust, and weathering of rocks [11–13]. But most of the elements enter and accumulate in the soil as a result of various types of human activities, such as (1) metallurgical mining and smelting (As, Cd, Pb, and Hg); (2) industry (As, Cd, Cr, Co, Cu, Hg, Ni, and Zn); (3) atmospheric deposition of transport emissions (As, Cd, Cr, Cu, Pb, Hg, and U); (4) agriculture (As, Cd, Cu, Pb, and Zn); and (5) waste disposal (eg, As, Cd, Cr, Cu, Pb, Hg, and Zn) [14–17].

The relative solubility of heavy metals under variability of environmental conditions causes differences in the relation of affinity to the environment and interaction with macroelements and toxicity in the composition of living organisms. Due to the low solubility, the role of trivalent or tetravalent cations Sn, Ce, Ga, Zr, and This practically reduced to zero. Of the remaining metals: Mo and Mn are important trace elements with low toxicity; Zn, Ni, Cu, V, Co, W and Cr are toxic elements of high and medium importance as trace elements and As, Ag, Sb, Cd, Hg, U and Pb have no useful function, but are considered toxins for cells [18, 19]

Lead is present in all components of the natural environment (0.0016% in the earth’s crust), participates in atmospheric-hydro spheric transport (most of it is deposited with dust, less - with atmospheric precipitation (less than 40%), enters plants from the soil and water, into the body of animals when consuming plants and water, into the human body - along with food, water and dust. At the same time, the main source of lead pollution for a long time was gasoline engines running on fuel that included tetraethyl lead, as well as thermal power plants (running on coal), mining, metallurgical and chemical industries. For example, lead was used to extinguish the reactor of the Chernobyl nuclear power plant, which then entered the air basin of the surrounding territories. So, in the air of industrial areas, the lead content can be 10,000 times higher than the natural level. Surface waters alone receive up to 300,000 tons of lead per year. Residents of industrialized countries have several times higher levels of lead in their bodies than residents of agrarian countries. Its deposition in bones, hair and liver increases [18, 19].

The mechanisms of the toxic effect of metals on the cells of organisms are generally well known, but it is difficult to define them as metal-specific concerning individual elements. This mechanism is competitive binding with proteins of essential and potentially toxic elements since metal ions stabilize and activate many proteins, being part of enzyme systems [19]. Some of the proteins have unoccupied SH groups that can interact with metal ions such as cadmium, lead, and mercury, leading to toxic effects. At the same time, the manifestation of the toxicity of metal ions in various organs and tissues of the body does not always correlate with the level of their accumulation [20, 21].

For microorganisms, the effectiveness of biological detoxification of metals through targeted microbial activity depends on factors such as culture age, cell shape, pH, contact time, and initial concentrations of metals in solution [22]. In this regard, several studies with actively growing cells have shown that some microorganisms can effectively remove heavy metal ions from the nutrient medium, especially when the concentration of metals is relatively low [23,24]. In particular, Mishra and Malik in 2012 [24] reported that the removal efficiency of a growing strain of Aspergillus lentils ranges from 34% for Ni^2 +^ and 71–78% for Cu^2 +^, Cr^3 +^, and Pb^2+^. Similar results were obtained when growing Zygosaccharomyces rouxii, in which the removal efficiency reached 94%, at the initial concentration of Cd ^2+^ it was less than 0.04 mm [25], while the removal efficiency when growing Beauveria bassiana was 61–75% for Cu ^2+^, Cr ^3+^, Cd ^2 +^, Zn^2+^ and Ni ^2+^ [26]. The above studies show that bioaccumulation by growing cells of microscopic fungi can provide a promising technology for the treatment of wastewater contaminated with heavy metals. In addition, direct use of actively growing cells simplifies system management and lowers operating costs by avoiding the need for a separate biomass production process [27].

Microscopic organisms have developed several strategies to overcome the inhibitory effects of toxic heavy metals. Thus, microbial strategies for metal detoxification include (1) elimination of metals by semi-permeable barriers; (2) active transport of metals away from the cell; (3) intracellular sequestration of metal by binding to protein; (4) extracellular sequestration; (5) enzymatic detoxification of a metal to a less toxic form; and (6) a decrease in the sensitivity of cellular targets to metal ions [28–30]. For any single metal or group of chemically bonded metals, one or more detoxification mechanisms can be used. However, the species of microorganisms can strongly influence the mechanisms of detoxification. For example, more microbes have specific stress resistance genes that can regulate resistance to toxic metals, and such regulatory genes are centrally located on plasmids or chromosomes. In general, the regulatory system for metal resistance is based on chromosomal genes, which are more complex than the plasmid system. On the other hand, systems encoded by plasmids usually interact with the mechanism of toxic ion efflux [31].

It has been reported that the determinants of plasmid-encoded resistance systems for toxic metal ions (such as copper) are inducible [32]. For example, the lead-resistant Enterococcus faecalis retained lead resistance even after the elimination of all plasmids [39], which indicates the location of signs of resistance to heavy metals in bacterial chromosomes [33].

Some Bacillus species present in saline soils produce exopolymers that give an adaptive advantage to bacteria to efficiently transport / remove heavy metals by biosorption [34].

Based on the analysis of the literature data, we set a goal: to study the mechanisms of biosorption of lead cations by representatives of bacteria of the genus Bacillus from the nutrient substrate when creating a massive cationic load on the environment and to assess the prospects of their use as microbial bioremediation in an in vitro model.

## EXPERIMENT DESIGN

### Research objects

The objects of the study were 6 strains of microorganisms of the genus *Bacillus* isolated from probiotic preparations: *B. subtilis* 534 (Sporobacterin, LLC Bakoren, Russia), *B. cereus* IP 5832 (Baktisubtil, Sanofi-Aventis, France), *B. subtilis* 10641 (Vetom 1), *B. licheniformis* 7038 (Vetom 2), *B. amyloliquefaciens* 10642 (Vetom 3), *B. amyloliquefaciens* 10643 (Vetom 4, Research Center, Russia, https://vetom.ru/).

### Research methods

To achieve our goal, the following methods were used in the work: the method of agar wells with the diffusion of cations into the thickness of the agar substrate combined with the method of serial dilutions, the nephelometric method for determining the relative optical density, atomic force microscopy, and atomic absorption spectrometry.

### Agar diffusion method

The method of diffusion in agar is the most indicative and easy-to-use method, which allows not only to assess the degree of dissociation of chemical compounds but also to determine the level of the biological effect of metal cations on the microorganisms under study. The study used the method of agar wells combined with the method of successive dilutions. The advantage of the method is the assessment of the effect of different concentrations of lead on the test organisms under study, which is in similar conditions (composition of the substrate, pH of the medium, temperature, etc.). This allows with a high level of confidence (the experiment was performed in more than 10-fold replication for each concentration) to assess the inhibitory effect of lead.

To implement the method, GRM agar was used as a nutrient medium, since all the studied microorganisms are chemo organoheterotrophs and this substrate provides their physiological needs. In the experiment, sterile aqueous solutions of Pb(NO_3_)_2_ and Pb(CH_3_COO)_2_ were used; the selection criterion for these chemical compounds is the high value of the dissociation constants in water, which makes it possible to create a high concentration of lead cations in the substrate (mobile forms) in a short period.

Daily cultures of microorganisms were plated on sterile nutrient media. Preliminarily, a series of dilutions of the studied chemical compounds were prepared at concentrations of 1 M, 0.5 M, 0.25 M, 0.125 M, 0.63 M, 0.031 M and 0.016 M. The choice of concentrations was determined by analyzing several published studies.

Wells with a diameter of 5 mm were made with a sterile microbiological punch according to a template in the thickness of the agar medium at a distance of 1.5 cm from the edge of the Petri dish and 3 cm between the wells in the amount of 7 on one dish. In the wells with a sterile automatic pipette, 30 μL of solutions containing various concentrations of metal were added (clockwise with a minimum concentration in the center of the dish). Cultures of bacterial cells in the presence of metal were incubated for 24 hours at 37 °C. The results were evaluated by measuring the diameter of the growth inhibition zones of the test organisms. This made it possible to determine the concentration of lead compounds that do not have a pronounced inhibitory effect on the studied microorganisms.

### Determination of the kinetics of growth of bacterial cultures

Determining the growth rate of a population of microorganisms in batch culture is a complex and time-consuming process. So, the counting of single cells is not only difficult to execute, but also does not exclude errors in their determination. Taking into account the fact that in the experiment it was required to determine not the number of cells, but the time the population reached a plateau at the maximum concentration, for which we used the nephelometric determination method in our work. The timing of the onset of the stationary growth phase was determined by measuring the optical density every 3 hours in daily cultures with introduced solutions of lead salts, relative to the control. For each series of repetitions of the study, a sterile culture medium GRM-broth was prepared and poured into sterile vials, 15 ml each. A suspension of microorganisms was preliminarily prepared, prepared according to the turbidity standard (0.5 McFarland) in saline. The vials were filled with 150 μl of the suspension. The background measurement was carried out immediately after the introduction of the suspension. The samples under study were incubated at a temperature of 37°C, 30 minutes before the measurement, they were placed on an incubator shaker to obtain a homogeneous suspension. Measurements were made in transparent plastic wells using a StatFax 303+ photometer (Awareness, USA).

### Determination of Pb content in solutions

The bioavailability was investigated by determining the lead content in the biomass of microorganisms by determining the difference in percentage between the control values of the element content in the nutrient substrate without adding a suspension of microorganisms and after their cultivation in the presence of lead salts using the method of atomic absorption spectrometry with flame atomization on an AAS instrument. 1 (Analytik Jena, Germany).

The samples under study were prepared by periodically cultivating the experimental test organisms in the presence of lead salts until the onset of the stationary growth phase. To accomplish this task, 300 ml of a liquid nutrient medium (GRM broth) prepared in solutions of lead salts and 3 ml of a suspension of microorganisms prepared according to the turbidity standard (0.5 McFarland) were added to vials with a volume of 400 ml, followed by cultivation until the onset of the stationary growth phase at a temperature of 37°C. Then the contents of the vials are centrifuged for 30 minutes at 3000 rpm to precipitate the microbial biomass. The supernatant was separated from the biomass content, the cell biomass was lysed by adding 5% KOH solution and kept for 20 minutes in a boiling water bath. Further, both the cell biomass and the supernatant were analyzed for lead content.

### Atomic force microscopy

To determine the spatial localization of the metal and to assess the morphometric characteristics of bacteria from the p. Bacillus atomic force microscopy was used. To accomplish this task, we used atomic force microscopy in this work using an original approach of sample preparation combined with the technique of agar wells and diffusion of the active substance into the thickness of the agar substrate.

Sample preparation consisted of fixing a freshly cleaved 5 × 5 mm mica plate to obtain a clean and atomically smooth surface, fixing it in the holder and contact of the plate with the surface of the nutrient medium on a Petri dish 1) in the growth inhibition zone near the well and 2) in the border zone between the zone of inhibition and the growth of the population of microorganisms on a dense nutrient medium.

Samples were scanned using an SMM-2000 atomic force microscope (ZAO PROTON-MIET, Russia) in contact mode using general-purpose cantilevers MSCT (Bruker, USA), rigidity 0.01 N / m and a nominal radius of curvature of the probe 10 nm. The quantitative morphometric analysis of the obtained images was carried out using the standard software of the microscope.

The data obtained during the experiments were subjected to statistical processing using primary methods for calculating elementary mathematical statistics: mean value, standard deviation, standard mathematical. Statistical processing of the results was carried out using the statistical software package Statistica 10.0. The obtained experimental data were processed by nonparametric methods with the calculation of the Student’s criterion, the differences were considered statistically significant p ≤ 0.05.

## RESULTS AND DISCUSSION

### Assessment of the toxicity of Pb (NO_3_)_2_ and Pb (CH_3_COO)_2_ about bacteria

The first stage of the research was to assess the inhibitory effect of lead salts on the test organisms under study. The data obtained during the study indicate the presence of a pronounced inhibitory effect on the growth of bacteria of concentrations of solutions of Pb(NO_3_)_2_ and Pb (CH_3_COO)_2_ from 1 M to 0.063 M about all studied bacteria (Figure 1). At the same time, the most stable about the studied chemical compounds in the maximum used concentration is the *B. subtilis* 10641 strains, the growth inhibition zone of which was 57.38% 1.57 times less than the analogous indicator of the most sensitive *B. amyloliquefaciens* 10643 strain (table 1)/ Concentrations (0.031 M and 0.016 M) of the investigated lead salts did not have a pronounced inhibitory effect.

**Figure 1.**
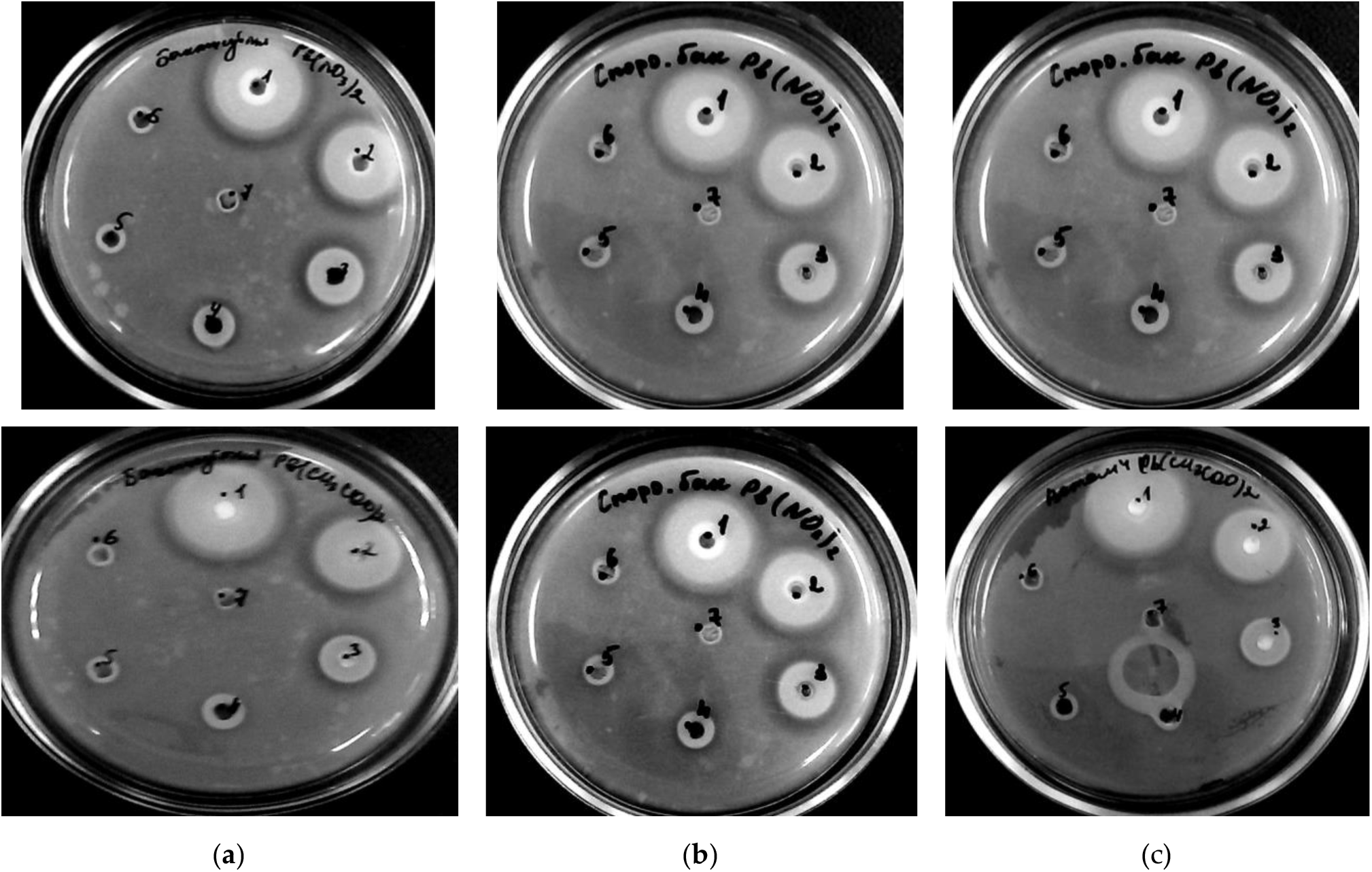
Effect of Pb(NO_3_)_2_ (top row) and Pb(CH_3_COO)_2_ (bottom row) on the growth of the studied microorganisms (in columns): (a) *B. cereus* IP 5832, (b) *B. subtilis* 534, (c) - *B. amyloliquefaciens* 10643 in concentrations from 1 M to 0.016 M (clockwise).

The zones of inhibition of the growth of the studied strains are insignificant differed in the presence of Pb(NO_3_)_2_ and Pb(CH_3_COO)_2_, with the most pronounced toxic effect observed at the maximum concentration of Pb(CH_3_COO)_2_. A high species sensitivity of *B. amyloliquefaciens* 10642 concerning high concentrations of lead nitrate and acetate has been experimentally established. Also, a relatively high level of resistance of *B. subtilis* 10641 about 1 M to 0.063 M solutions was noted, which probably indicates the presence of individual characteristics of this microorganism to show maximum resistance to a high cationic load of lead.

### Evaluation of the effect of lead on the growth characteristics of the studied strains in batch culture

The next stage of our research was to assess the effect of lead nitrate and acetate on the growth of tested microorganisms under conditions of periodic cultivation in a liquid substrate. Experimental studies have shown that the concentration of Pb(NO_3_)_2_ and Pb(CH_3_COO)_2_ 0.031 M did not have a pronounced inhibitory effect against the selected bacterial strains, therefore the effect of lead on the growth of the population of each bacterial strain was studied by the nephelometric method (Figure 2).

**Figure 2.**
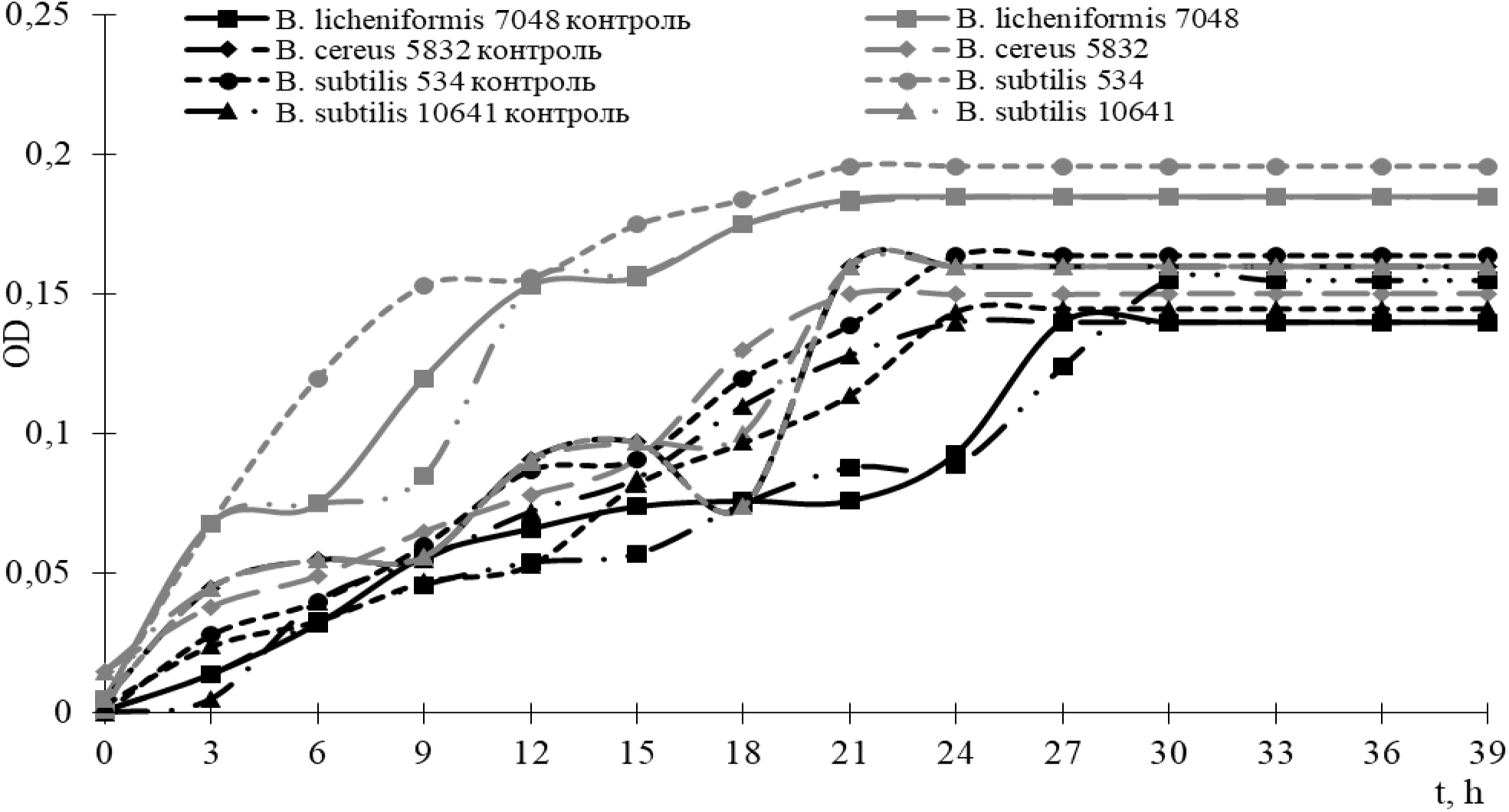
Effect of Pb (NO3) 2 at a concentration of 0.031 M on the growth dynamics of the studied microorganisms

The stepwise change in the density of cells of all studied strains in the phase of exponential growth is due to the restructuring of the enzyme systems of microorganisms as the substrate is depleted. It should be noted the general tendency of stabilization of indicators of an increase in bacterial mass by the 24th hour of the experiment and the onset of the stationary growth phase.

Comparative analysis of the curves indicates that the populations of *B. subtilis* 534, *B. cereus* 5832, and *B. licheniformis* 7038 reached maximum density under conditions of increased cationic loading. Lead does not participate in the metabolic processes of microorganisms, therefore, the growth of cell density cannot be justified by the stimulating effect of the element. Explain this phenomenon by the high sorption characteristics of the surface structures of the cells of the presented strains with the subsequent formation of conglomerates.

The rest of the strains did not have significant differences in the values of cell density in comparison with the control values against the background of the presence of lead salt in the nutrient substrate.

### Determination of spatial localization on the surface of bacterial cells and its effect of lead on morpho-physiological characteristics

The use of atomic force microscopy made it possible to conduct a visual assessment of the toxicity level of Pb(NO_3_)_2_ and Pb(CH_3_COO)_2_ and the degree of influence of lead cations on the morphological characteristics of bacterial cells (Figure 3). Determination of spatial localization on the surface of bacterial cells and its effect of lead on morpho-physiological characteristics.

**Figure 3.**
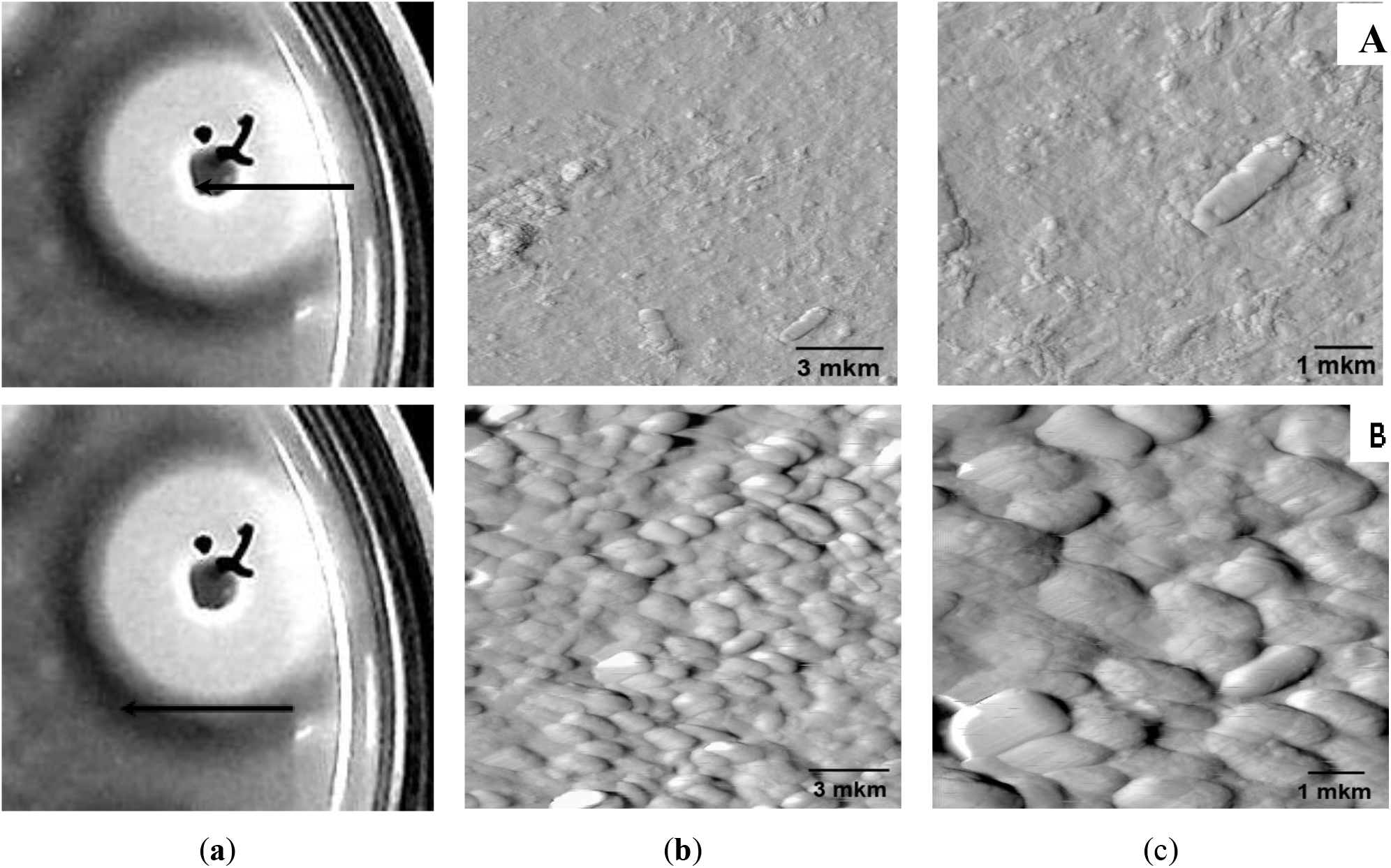
Evaluation of the effect of lead cations on the example of B. subtilis 534 using atomic force microscopy: A - sampling zone for AFM in the growth inhibition zone, B - sampling for AFM in the boundary growth zone of the studied microorganisms

The use of atomic force microscopy made it possible to conduct a visual assessment of the toxicity level of Pb(NO_3_)_2_ and Pb(CH_3_COO)_2_ and the degree of influence of lead cations on the morphological characteristics of bacterial cells (Figure 3).

The experimental data obtained indicate a high level of lead diffusion into the structure of the agar nutrient medium, while a high level of interaction of the xenobiotic element with the components of the substrate with its maximum deposition on the surface of the medium should be noted (Figure 3 A). It is very difficult to assess the level of viability of the studied strains in the zone of element localization since separately located cells with the maximum level of interaction with lead cations without changing their morphological characteristics are visually recorded. At the same time, in the studied areas, there are no sporulating vegetative forms and spores of the studied microorganisms.

Hypothetically, this phenomenon can be explained as follows: a high level of cationic load of a xenobiotic on a substrate provokes the adaptive and physiological mechanisms of individual representatives of population groups, but at the same time blocks their metabolic processes and mechanisms of sporulation, as a result of which the bacterial cell loses its ability to divide and form resting forms.

As the radial distance from the hole is formed, a zone is formed characterized by the absence of a visible effect of lead cations and the absence of growth in the population of the studied microorganisms, which in turn allowed us to determine the next boundary zone for the selection of experimental samples (Figure 3B). A visual assessment of the degree of influence of subinhibitory concentrations of lead on the test organisms under study indicates a high loss of sorption of the element on the cell surface, while, in contrast to the sample of the growth inhibition zone in this area, active growth of vegetative forms of bacterial strains and sporulation are recorded, which in turn indicates a negative the influence of xenobiotics on metabolic processes and, as a consequence, the inclusion of physiological-adaptive mechanisms of resistance to the effects of unfavorable environmental factors. The concentration of lead cations forming in this zone has a significant effect on the morphological characteristics of the studied strains, manifested in the form of shortened forms of bacterial cells and the absence of a characteristic arrangement of cells relative to each other (streptobacilli).

In our opinion, the detoxification mechanism is due to the formation of biologically inactive forms of metal on the surface of bacterial cells, which in turn ensures the safety of the population.

### Biosorption characteristics of microorganisms

As part of confirming the hypothesis of the mechanism of detoxification of an element associated with sorption on the cell surface, we carried out a study aimed at assessing the bioaccumulative characteristics of the studied strains of microorganisms about lead cations (Figure 4), for which we used the method of atomic absorption spectrometry.

**Figure 4.**
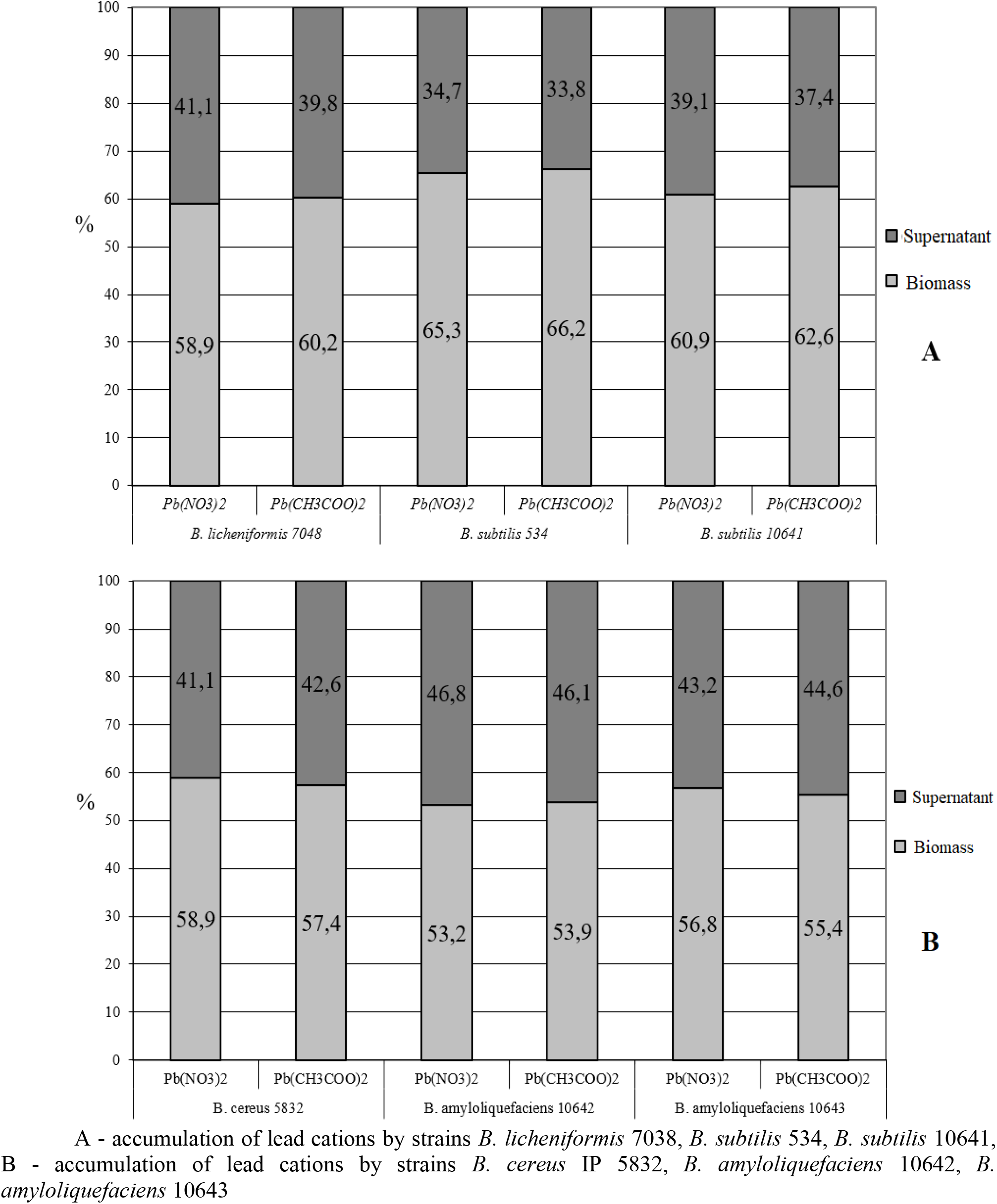
Assessment of the sorption characteristics of bacteria of the genus Bacillus of lead cations from the substrate with the addition of Pb(NO_3_)_2_ and Pb(CH_3_COO)_2_

Under the experimental conditions, Pb(NO_3_)_2_ and Pb(CH_3_COO)_2_ at a dose of 0.031 M were added to the liquid substrate before the introduction of the tested microorganisms, followed by incubation of bacterial strains in the presence of a xenobiotic element for 30 hours at a temperature of 37 ° C, which was determined earlier. The results were accounted for by the difference between the doses of the introduced element and its determination in the biomass and supernatant.

Analysis of the experimental data obtained indicates a pronounced biosorption activity of all studied strains about lead cations from the substrate. At the same time, no significant differences were found in the tested samples both in the presence of Pb(NO_3_)_2_ and Pb(CH_3_COO)_2_, respectively. The heterogeneity of the results obtained (Figure 4) does not allow us to reveal a general pattern on the degree of influence of the anionic component on the level of sorption. However, in the course of the study, we established the species and strain accumulating activity of individual representatives of microorganisms of the genus Bacillus.

Thus, the maximum values of sorption were recorded in representatives of the species *B. subtilis*, the percentage of biosorption of which at the stage of reaching the stationary growth phase averaged 65.3% for B. subtilis 534 and 66.2% in the presence of Pb(NO_3_)_2_ and Pb(CH_3_COO)_2_, respectively. For the *B. subtilis* 10641 strain, these values were 60.9% and 62.6%, respectively. Of all the studied microorganisms, representatives of the species *B. amyloliquefaciens* have the minimum accumulating characteristics (Figure 4). However, it should be noted that the use of all bacterial strains is promising in the implementation of projects for the use of bacterial strains of the genus *Bacillus* as bioremediation of biologically active forms of lead in ecological systems of various levels of the organization.

The combination of the methods used made it possible to fully assess the degree of toxicity and the effect on the growth of lead cations on the studied microorganisms in a model experiment. The use of atomic absorption spectrophotometry and atomic force microscopy made it possible to assess the degree of biosorption of a xenobiotic element from a substrate with the localization of inactive forms of lead.

## CONCLUSIONS

The experimental data obtained indicate the presence of a directly proportional relationship between the level of toxicity, the degree of influence on growth and the sorption characteristics of the studied strains.

The high resistance of the *B. amyloliquefaciens* strains to Pb(NO_3_)_2_ and Pb(CH_3_COO)_2_ and the sensitivity of the *B. subtilis* 10641 strains, which are most pronounced in the maximum concentration and level with a decrease in the content of metal ions, are shown. At the same time, no significant differences from the composition of the anionic component of the test compound were found.

The degree of influence of the lead load on the growth characteristics of the studied strains in batch culture is determined, which is expressed in the completion of a dynamic increase in biomass by the 24th hour of the experiment using a minimum concentration of 0.031 M, which does not have a pronounced inhibitory effect on the experimental bacterial strains.

The data obtained were used to study the biosorption characteristics of the tested microorganisms about lead cations and to determine the place of their localization and the degree of influence on the morpho-physiological characteristics of the strains. As a result, a high degree of biosorption of all tested microorganisms was established with the distribution of indicators in the presence of Pb(NO_3_)_2_ and Pb(CH_3_COO)_2_ in the substrate for *B. subtilis* 534 - 65.3% and 66.2%, *B. subtilis* 10641 - 60, 9% and 62.6%, *B. licheniformis* 7038 - 58.9% and 60.2%, *B. cereus* IP 5832 - 58.9% and 57.4%, *B. amyloliquefaciens* 10643 - 56.8% and 55.4%, *B. amyloliquefaciens* 10642 - 53.2% and 53.9%, respectively.

It was experimentally established that there is no pronounced effect of the anionic component in the structure of chemical compounds on the level of biotoxicity and sorption of lead cations.

Investigation of the detoxification mechanisms of an element hypothetically associated with the sorption of an element on the surface of bacterial fouling in a biologically inactive form, which is confirmed by the results obtained using AFM, which coincide with the literature data [35]. For example, it was experimentally established by Schroeter et al. That isolates of *B. licheniformis, B. cereus* and *B. subtilis* have the maximum surface adsorption of lead as determined by atomic absorption spectrometry and additionally by scanning electron microscopy. The binding mechanism includes a combination of ion exchange, chelation, and sometimes reductive reactions, accompanied by the deposition of metallic Pb on the cell wall material. It is believed that in the course of ion-exchange processes, the cations of Ca, Mg, H, Na, K in the material of the cell walls are replaced by heavy metals [36]. In works [37, 38] it was found that biosorption of lead modifies the carboxyl-, hydroxyl-, amino-, sulfhydryl and sulfate groups of peptidoglycans, where ions of other metals cannot compete, offering it a greater affinity.

Summarizing the above, it should be noted that there is a high level of interest in the study of the biosorption characteristics of microorganisms for the removal of heavy metals with the prospect of their use as bioremediation in ecological systems of various levels of the organization, that is, to solve the problems of cleaning the natural environment.

## Acknowledgment

The research was supported by the Ministry of Science and Higher Education by the state assignment for Ural State Mining University No. 0833-2020-0008 ‘Development and environmental and economic substantiation of the technology for reclamation of land disturbed by the mining and metallurgical complex based on reclamation materials and fertilizers of a new type’. We obtain the scientific results with the staff of Center for the collective use by using funds of the Center for the collective use of scientific equipment of the Federal Scientific Center of biological systems and agricultural technologies of RAS as well (No Ross RU.0001.21 PF59, the Unified Russian Register of Centers for Collective Use - http://www.ckp-rf.ru/ckp/77384).

