## Supplementary material for "Estimation of the biosorption potential of certain representatives of the genus Bacillus in interaction with lead cation *in vitro*": Table 1

Table 1. Diameter of the zone of suppression (mm) of representatives of the genus *Bacillus* with various concentrations of lead salts

| Test strains | The concentration of the investigated substance, M | | | | | | |
|---|---|---|---|---|---|---|---|
| | 1 | 0,5 | 0,25 | 0,125 | 0,063 | 0,031 | 0,016 |
| **$Pb(NO_3)_2$** | | | | | | | |
| *B. licheniformis* 7048 | 31,0±1,0 | 27,3±0,3 | 20,3±0,7 | 15,7±0,3 | 7,0±1,5 | 0 | 0 |
| *B. cereus* 5832 | 30,3±0,3 | 25,3±0,7 | 19,3±0,3 | 10,7±1,2 | 7,3±0,3 | 0 | 0 |
| *B. subtilis* 534 | 30,0±0,01 | 26,0±0,6 | 17,0±0,6 | 9,7±0,3 | 7,1±0,6 | 0 | 0 |
| *B. subtilis* 10641 | 23,7±0,3 | 18,7±0,3 | 14,7±0,3 | 9,7±0,3 | 6,7±0,3 | 0 | 0 |
| *B. amyloliquefaciens* 10642 | 36,0±0,6 | 27,7±1,2 | 24,7±2,2 | 10,7±0,3 | 7,3±0,3 | 0 | 0 |
| *B. amyloliquefaciens 10643* | 37,3±1,5 | 30,3±0,9 | 20,0±2,9 | 10,7±0,3 | 6,7±0,3 | 0 | 0 |
| **$Pb(CH_3COO)_2$** | | | | | | | |
| *B. licheniformis* 7048 | 32,0±0,01 | 27,0±1,0 | 20,7±1,7 | 12,3±0,3 | 9,1±1,9 | 0 | 0 |
| *B. cereus* 5832 | 32,3±0,3 | 22,3±1,5 | 14,0±1,4 | 9,7±0,3 | 7,7±0,3 | 0 | 0 |
| *B. subtilis* 534 | 32,0±0,6 | 25,3±0,3 | 13,3±0,9 | 10,3±0,3 | 6,4±0,3 | 0 | 0 |
| *B. subtilis* 10641 | 23,3±0,3 | 19,3±0,3 | 13,0±0,6 | 9,7±0,3 | 6,7±0,3 | 0 | 0 |
| *B. amyloliquefaciens* 10642 | 38,7±0,3 | 30,3±0,9 | 23,3±0,9 | 10,3±0,3 | 7,0±0,6 | 0 | 0 |
| *B. amyloliquefaciens 10643* | 31,7±1,7 | 22,3±0,3 | 13,3±0,3 | 8,7±0,9 | 5,7±0,7 | 0 | 0 |
